# Evaluating Minocycline as a Neuroprotective Agent in an Aged Female Rat MCAO Stroke Model

**DOI:** 10.1101/2023.06.13.544795

**Authors:** Liza Gutierrez, Vincent M. Tutino, Adnan H. Siddiqui

**Affiliations:** Canon Stroke and Vascular Research Center; University at Buffalo, Buffalo, NY, USA; Department of Pathology and Anatomical Sciences, University at Buffalo, Buffalo, NY, USA; Department of Neurosurgery, University at Buffalo, Buffalo, NY, USA

## Abstract

The middle cerebral artery occlusion (MCAO) suture model is widely accepted ischemic stroke model. However, researchers routinely use young male rats, ignoring that stroke risk is increased in older, post-menopausal women. To this end, we implemented (120-minute) transient- and permanent-occlusion MCAO models in female retired-breeder rats, examining the endpoint across 1-30 days to identify the optimal time course for the model. We found that in both groups the physical infarct (measured by triphenyltetrazolium chloride -TTC staining), which is present initially, was not detectable 30 days post-MCAO (even if some neurologic symptoms persist). Across shorter time-points (namely 24 hours and 7 days) we found that neurologic scores generally reach a plateau/maximum at ∽7 days, then infarct size gradually decreases over time for rats receiving a permanent MCAO. Across 3 transient occlusion times (60 minutes, 90 minutes, and 120 minutes), the longest gave the most robust result. Overall, the permeant and 120-minute transient MCAO evaluated at 7 days was optimal. Using these two models, we evaluated the neuroprotective qualities of the antibiotic, minocycline. We found that those in the treatment groups experienced a greater improvement in neurologic scores and a larger decrease in infarct size after daily treatment for seven days. This improvement was more prominent in the transiently occluded treatment group than in the permanently occluded group.

## Introduction

Ischemic stroke is a life-threatening condition that typically occurs when a blood vessel that carries oxygen and other nutrients to the brain tissue is blocked by a blood clot. Every year approximately 15 million strokes occur worldwide.^1^ In the United States, approximately 800,000 people experience a stroke every year, which equates to one stroke every 40 seconds.^2^ Despite advancements in treatment and interventional strategies, such as the administration of recombinant tissue plasminogen activator or mechanical thrombectomy techniques (like stent retrievers or aspiration catheters), the outcomes following a stroke remain poor. There are still high rates of long-term disability following stroke, high rates of stroke recurrence (secondary stroke),^3^ and cases of cryptogenic stroke where diagnosis of etiology is not possible.

Towards improving stroke outcomes, finding better pharmacological treatments, and improving secondary prevention, researchers continue to experimentally study stroke in animal models. The most popular *in vivo* model of stroke remains the middle cerebral artery occlusion (MCAO) suture model. Although MCAO is widely-accepted as a reliable and repeatable animal model of stroke, there are several challenges that the model has yet to address. In particular, this model is pronominally implemented in young, male rodents. An increasing prevalence for stroke in women, particularly those later in life,^2,4^ and a lack of corresponding female animal data, suggest that implementing MCAO in older, female rats should be considered. Moreover, most models examine infarct size shortly (less than three days) after MCAO. There have been no published reports in the literature that have investigated the duration and timeline for infarct analysis following MCAO.

To this end, we sought to implement the MCAO model in female, retired breeder rats, and sacrifice them at multiple time points to determine the most appropriate end point for analysis. Furthermore, because stroke can present with varying ischemic timelines, we also sought to induce permanent occlusions as well as transient occlusions across different timepoints. The best model (for optimal infarct size and survivability) was then employed in a preliminary study to test the effect of minocycline on stroke severity. Minocycline is a synthetic, second generation tetracycline, has been shown to exhibit neuroprotective qualities in animal models of multiple sclerosis, Parkinson’s disease,^5^ and Huntington’s disease.^6,7^ Although its mechanism of action remains unknown, minocycline may have anti-inflammatory effects^8^ that may reduce microglial activation,^9,10^ reduce matrix metalloproteinase activity,^11^ or inhibit cell death by blocking activated caspase 3 formation.^12^

## Methods

### MCAO Models

Female retired breeder Sprague Dawley rats (Harlan) weighing 280-320g were used for all surgeries. Cerebral ischemia, both transient and permanent, was induced by intraluminal suture occlusion of the MCA.^13^ A 30 mm 4-0 monfilament silk suture with a silicone coating length of 2-3 mm and diameter of 0.37-0.39 mm (Doccol) was used to occlude the MCA origin. Access to the MCA origin was obtained by transecting the external carotid artery (ECA), temporarily placing an artery clip on the common carotid artery (CCA), and using the external artery stump to pass the suture into the internal carotid artery (ICA) toward the MCA. The MCA origin is known to be approximately 2.0 cm from the origin of the ICA. Once the suture was inserted the appropriate distance, it was tied into place with a 4-0 silk suture ligation around the ECA stump and the artery clip around the CCA was removed. The suture was kept in place for the permanent occlusion models and removed after the desired time for the transient occlusion models. The incision was then closed and the rat was recovered from anesthesia. At the time of sacrifice, rats were placed under anesthesia with 5% isoflurane, and then euthanized by intraperitoneal administration of 100mg/kg of sodium pentobarbital. All care and procedures were performed in accordance with institutional guidelines as approved by the local Institutional Animal Care and Use Committee.

All rats were evaluated six hours post-operatively, and then daily until sacrifice for weight and hydration, and non-invasively graded using Menzies scoring to estimate stroke severity (see Table 1). Menzies et al.^14^ details the score (0-4) as: 0) normal animal extends both forelimbs symmetrically toward the floor, 1) contralateral forelimb is consistently flexed during suspension, 2) decreased grip of the contralateral forelimb when pulled by the tail, 3) monodirectional circling toward the paretic side, when a slight jerk of the tail is given, and 4) consistent spontaneous contralateral circling. Higher scores are cumulative of the responses present in lower scores. Rats showing no deficit 24 hours after the MCAO procedure were excluded from the study.

**Table 1.** Menzies neurologic scoring.

| **Score** | **Evaluation Metric** |
| --- | --- |
| 0 | No apparent deficits |
| 1 | Contralateral forelimb flexion |
| 2 | Decreased grip of the contralateral forelimb when tail pulled |
| 3 | Spontaneous movement in all directions; contralateral circling, only if pulled by tail |
| 4 | Spontaneous contralateral circling |

### Brain TTC Staining

TTC staining is a widely accepted technique to differentiate between metabolically active and inactive tissue.^15^ TTC (GFS Chemicals) dissolved in phosphate buffered saline (PBS) at a 0.1% concentration was warmed to 37°C in a hot water bath. Immediately following euthanasia, the brain was harvested and placed into a matrix filled with chilled PBS and sliced into seven 2 mm coronal sections, excluding the cerebellum. Sections were placed into warmed TTC solution in a 12-well uncoated cell culture plate, wrapped in foil, and incubated in a 37°C water bath for approximately 10 minutes. Slices were turned over once to allow equal staining and monitored every 2 minutes until staining was complete. Slices were then transferred to other wells containing 10% formalin and allowed to fix for at least 4 hours in the dark before being scanned. After 4-24 hours of fixation, slices were arranged onto a glass slide and scanned using an HP color desktop scanner.

To determine the degree of infarct in rats, TCC staining was quantified.^16^ Metabolically active areas of the brain stain red when dehydrogenases in healthy tissue reduce TCC into a red compound, triphenylformazan. Inactive areas of the brain that do not metabolize TCC and are represented by white discoloration of the tissue. Image J was used to measure the red areas of healthy tissue and white areas of infarction in each of the 7 brain sections for each specimen. Three independent measurements of cerebral infarction area were measured for each brain slice, totaling 21 infarction measurements for each of the 7 animals. Each measurement was then multiplied by section thickness and divided by the total brain volume of each section to calculate percentage of infarction, as detailed below.

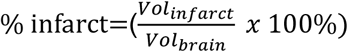

### Model Time-Course Optimization

We sought to to optimize the MCAO protocol in terms of the severity of infarct and length of treatment time. To do this we implemented permanent occlusions in n=22 rats, 60-minute occlusions in n=2 rats, 90-minute occlusions in n=10 rats, and 120-minute occlusions in n=11 rats. In each group, we then investigated the infarct size at different time points, namely, 24 hours, 7 days, and 30 days (which was done because the 30 day this time point more closely mimics human treatment protocols). After evaluating the data from all timepoints, and in order to balance the severity of infarct with length of treatment time, a the permanent and the 120- minute transient MCAO models with a 7-day analysis time-point were chosen to repeat and use for later treatment groups described in the next section.

### Minocycline Treatment

We performed experiments to test if treatment with minocycline could decrease the percent infarction following stroke in our rat models. A total of n=10 rats (n=10 permanent MCAO and n=10 120-minute transient MCAO) received minocycline and n=10 rats (n=10 permanent MCAO and n=10 120-minute transient MCAO) received a vehicle control.

Minocycline powder (Sigma-Aldrich) was dissolved in sterile saline at a concentration of 13.5 mg/ml and filtered through a 0.40 μg filter prior to delivery. The minocycline solution was stored covered at 4°C for no more than 5 days before use, per supplier recommendations. The dosing regimen we followed was based on previous studies.^8,10,11,17,18^ Minocycline was delivered intraperitoneally to treatment animals in two doses of 45 mg/kg, the first being immediately post-operatively and the second 6 hours later. These animals received daily intraperitoneal injections of 22.5 mg/kg minocycline until sacrifice at 7 days. “Vehicle control” animals received intraperitoneal injections of the same volume of sterile saline.

### Statistical Analysis

All statistical analysis was completed in Microsoft Excel. We used an α=0.05 to determine statistical significance. Comparison of mean infarct morphometrics between groups was completed using a Student’s t-test.

## Results

### Model Optimization and Parameter Selection

In tailoring the model, we subjected rats to either permanent occlusion (n=22) or transient occlusion of 60 minutes (n=2), 90 minutes (n=10), or 120 minutes (n=11) (Table 2). While control animals sacrificed at 24 hours showed significant infarcts after occlusion times of 90 and 120 minutes or permanent occlusion, a 60-minute occlusion time resulted in 50% mortality (one died) and no infarct in the surviving animal (n=1). Due to this variability, no further animals were completed for this occlusion time. We analyzed infarcts at 24 hours, 7 days, and 30 days after the procedure. Strikingly, the permanent (as well as the 90-minute transient) MCAO had no visible infarct at 30 days. Figure 1A demonstrates this on TTC staining, which is quantifies as percent infarct in Figure 1B. Neurologic scoring also showed a gradual downward trend over the course of 30 days in the permanent MCAO group and did so in an almost linear fashion, with a 93.4% correlation (Figure 2). This suggests that over time, stroke injury decreases in aged female rats.

**Table 2.** Numbers of animals used to investigate occlusion time course.

| **Occlusion** | **24 Hours** | **2 Days** | **7 Days** | **10 Days** | **30 Days** |
| --- | --- | --- | --- | --- | --- |
| *Permanent* | n=5 | n=2 | n=5 | n=2 | n=8 |
| *60-minute* | n=1* | - | - | - | - |
| *90-minute* | n=2 | - | n=2 | - | n=6 |
| *120-minute* | n=5 | n=1 | n=5 | - | - |

\* Two rats were completed, one rat expired post-operatively

**Figure 1:**
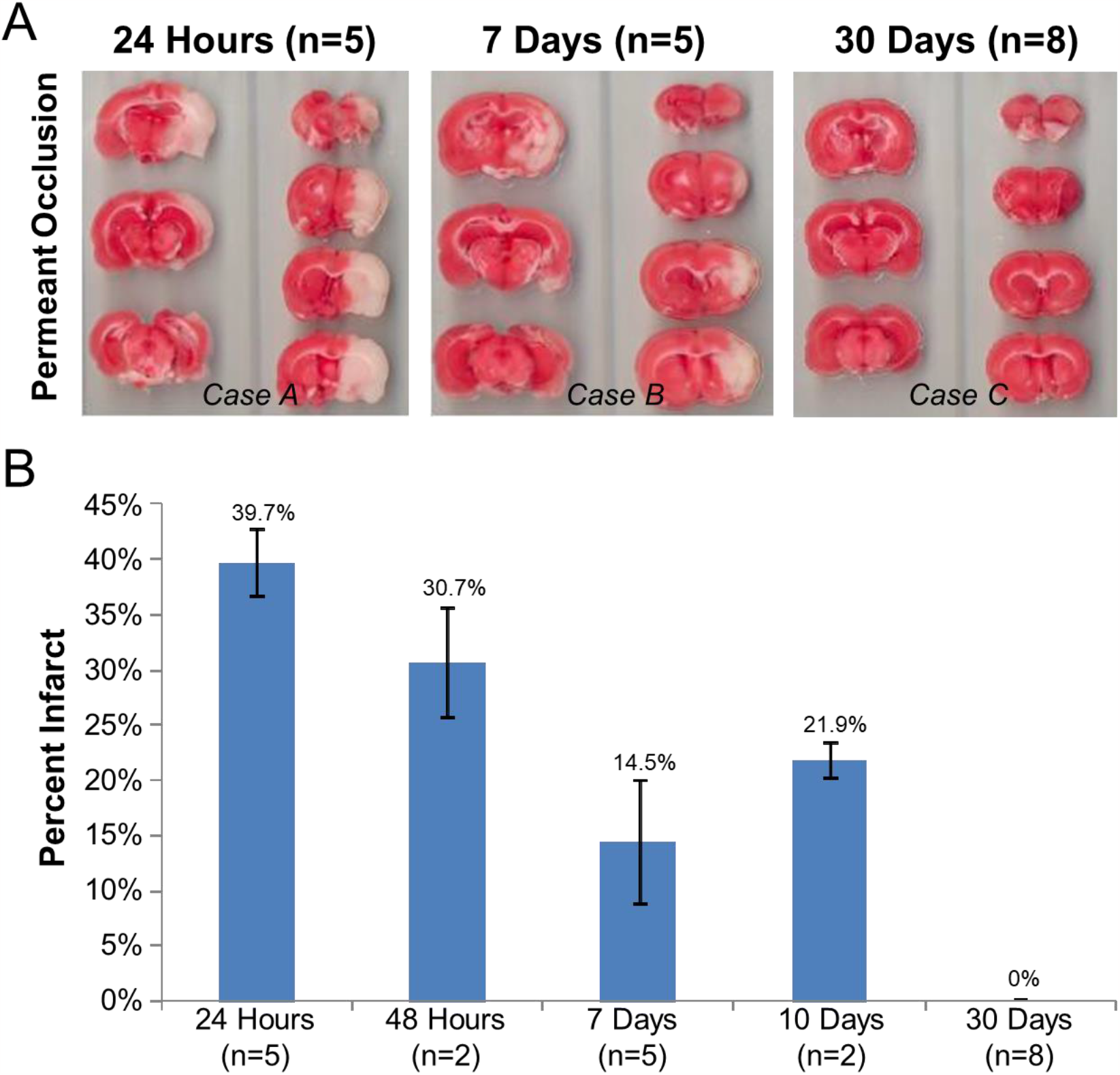
Permanent MCAO infarct size over 30 days. **A)**. Areas of infarction appear white, while healthy tissue is stained pink. A representative case at each time point was chosen. **B)**. Average percent infarct of permanent MCAO control rats at all time points collected with standard error bars.

**Figure 2:**
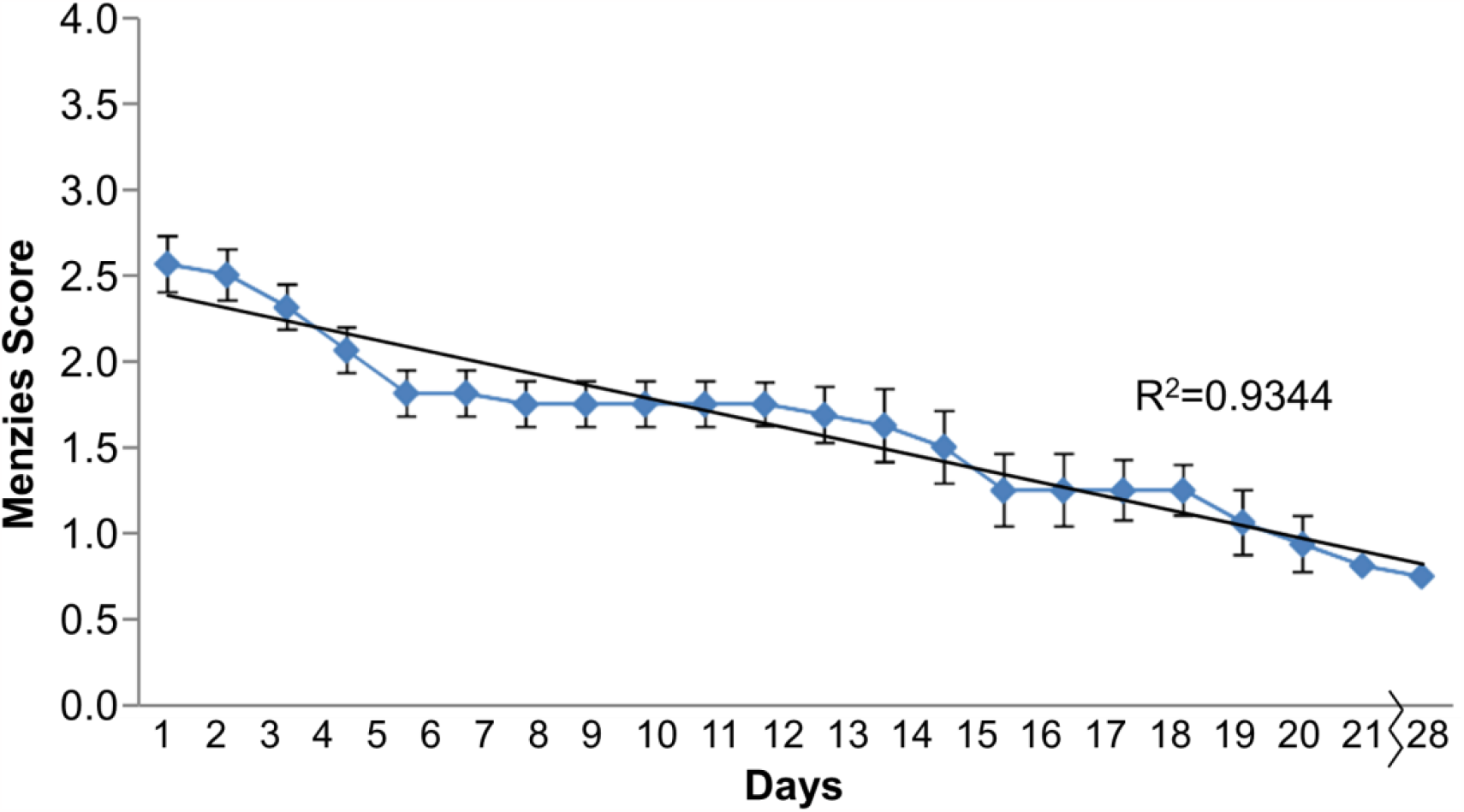
Neurological scoring over 30 days. Average Menzies scores of the 30-day permanent MCAO control group with standard error bars and correlation.

While unplanned, n=2 rats that had been part of a permanent MCAO group required euthanasia at a 2-day time point. We noted the presence of a large infarct (25.8% and 35.7% direct infarct) in these animals (see Supplemental Figure 1, Left). There were also n=2 permeant MCAO rats that required euthanasia at 10 days. Both were found to have unusually large infarcts (23.5% and 20.2%) for that length of time post-operatively (larger than the average infarct size of the 7-day permanent MCAO group) (see Supplemental Figure 1, Right). The severity of symptoms forced the early removal of these rats from the 30-day group.

After the 7-day end-point was established, we analyzed data for occlusion time for the transient group. As shown in Figure 3A, we found that the found that even at 7 days, the apparent infarct was too far diminished in the 90-minute transient MCAO group. Therefore, it was decided to extend the length of occlusion to 120 minutes (Figure 3B). Increasing the occlusion time resulted in a visible infarct remaining through the 7-day end point. Thus, for the transient MCAO procedure an occlusion time of 120 minutes was recommended with a sacrifice time point of 7 days for analyzing changes in infarct.

**Figure 3.**
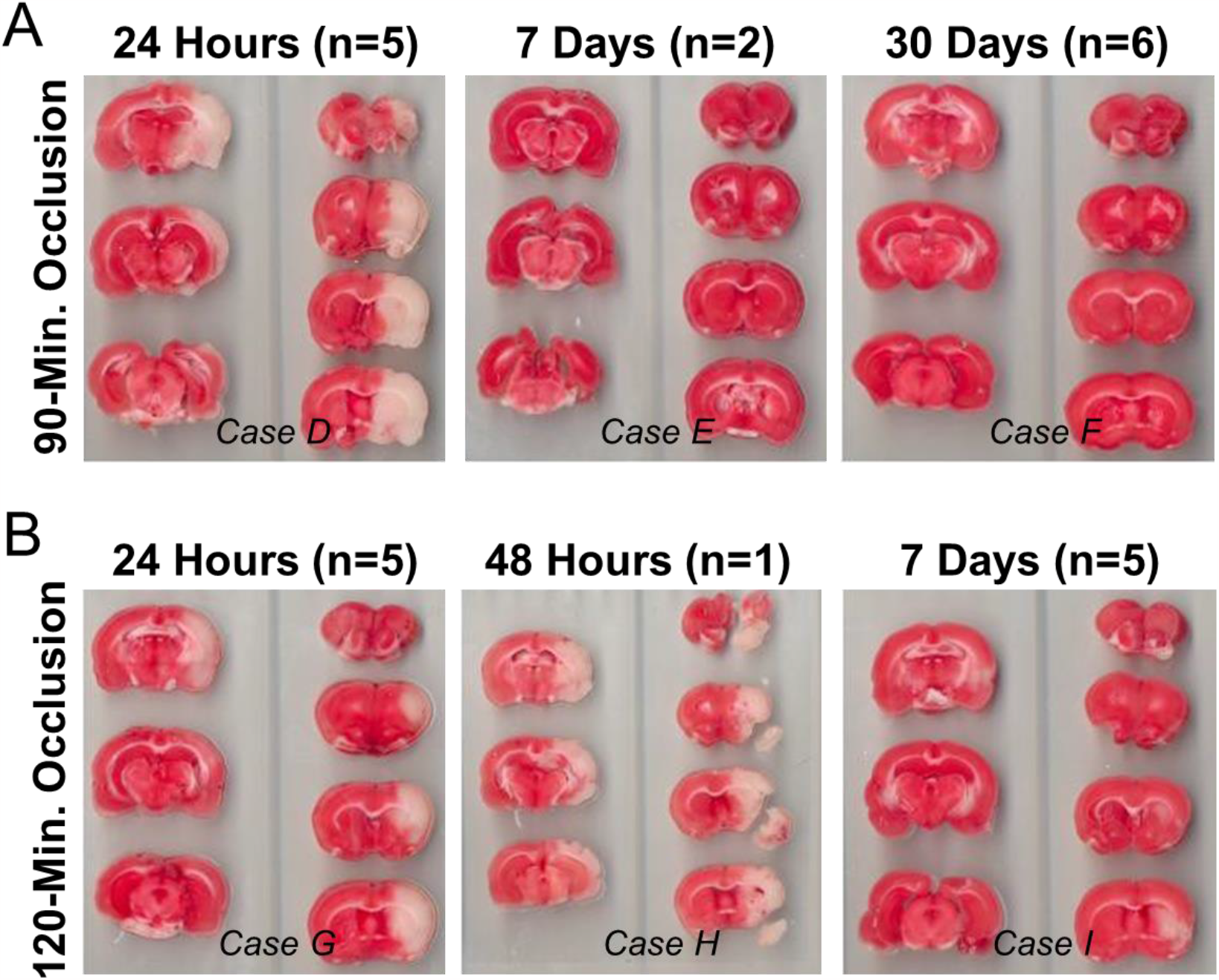
Thee 90-minute and 120-minute occlusion cases. **A)**. 90-minute transient MCAO infarct size over 30 days. Areas of infarction appear white, while healthy tissue is stained pink. A representative sample for each time point was chosen. **B)**. 120-minute transient MCAO infarct size over 7 days. Areas of infarction appear white, while healthy tissue is stained pink. In groups with more than one sample, a representative image for each time point was chosen.

### Minocycline Treatment Reduces Infarct Size at 7 Days

We used a group of n=20 rats (see Table 3 for groups and numbers of animals) for a preliminary study investigating the effect of minocycline on infarct. All experimental animals that received permanent MCAO procedures but were not treated with drugs exhibited infarct development. After a seven-day course of minocycline, the average infarct size following permanent MCAO decreased compared to control animals receiving only a saline placebo (Figure 4A). Though not statistically significant, this data also corresponded with a slight improvement in Menzies scores in treatment animals when compared with those in the control group (Figure 4B). These experiments were also run in the transient MCAO model. In animals receiving a 120-minute transient MCAO, a trend showing decreased infarct size was also apparent in the minocycline treatment group when compared to the no-treatment controls, although variability and a small sample size likely prevented statistical significance from being achieved (Figure 4C). This data also corresponded with an improvement in Menzies score in the treatment group versus control animals, showing a greater improvement in neurologic scores over the treatment period (Figure 4D).

**Table 3.** Numbers of animals used in the minocycline study.

| **Occlusion** | **Treatment** | **7 Days** |
| --- | --- | --- |
| *Permanent* | Minocycline | n=5 |
| *Permanent* | Vehicle Control | n=5 |
| *120-minute* | Minocycline | n=5 |
| *120-minute* | Vehicle Control | n=5 |

**Figure 4:**
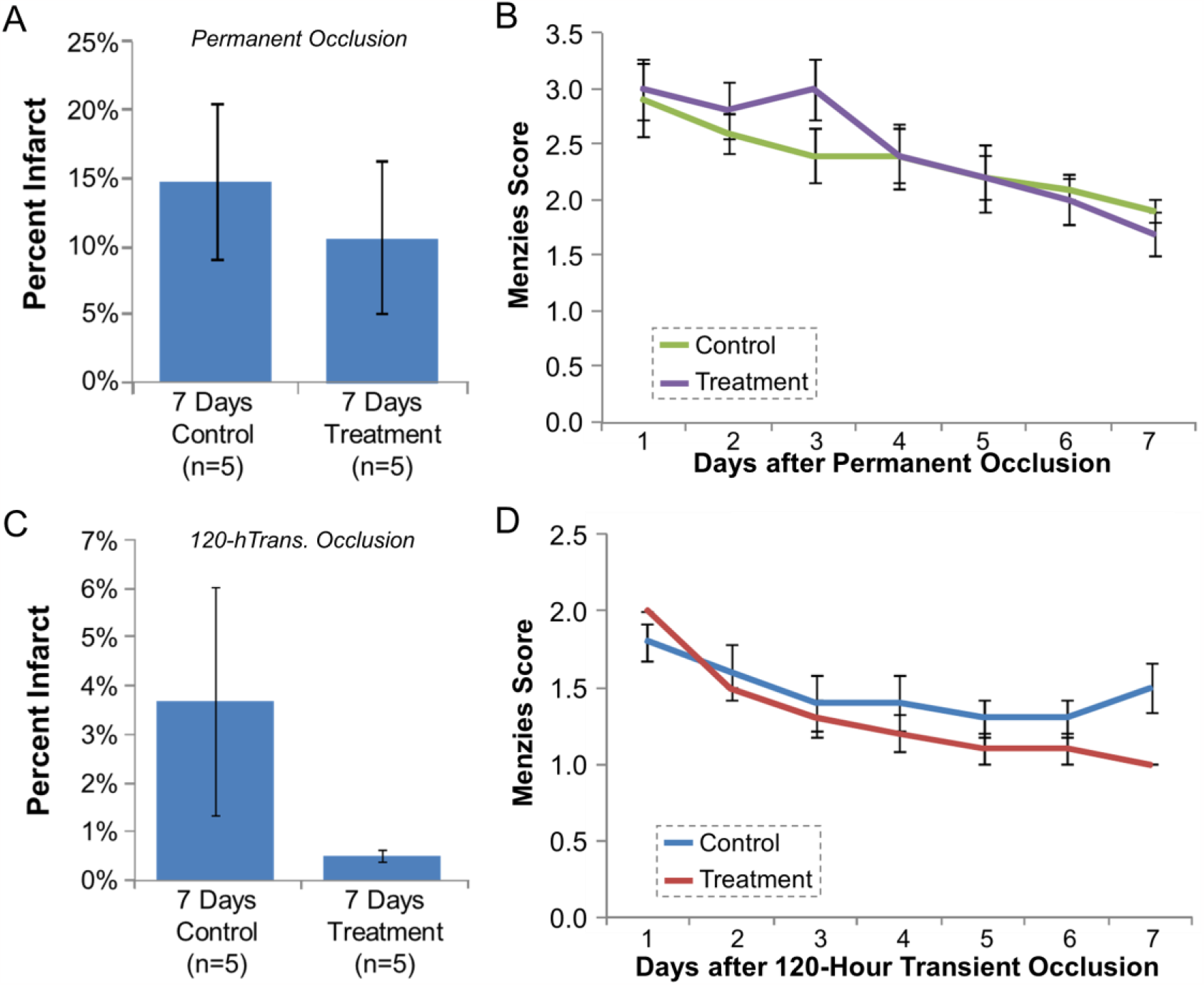
Percent infarct and neurologic scoring for minocycline treated strokes and controls. **A)**. Average percentage of direct infarct for permanent MCAO control and treatment groups with standard error bars. **B)**. Average Menzies score for permanent MCAO control and treatment groups with standard error bars (n=5 for each group). **C)**. Average percentage of direct infarct for 120-minute transient MCAO control and treatment groups with standard error bars. **D)**. Average Menzies score for 120-minute transient MCAO control and treatment groups with standard error bars (n=5 for each group).

## Discussion

While the MCAO rat suture model is a widely accepted model, this is the first study to evaluate longer time points in retired breeder female rats and also the first to to examine infarct sizes and track neurologic scores over a 30-day period. This information is particularly useful for pharmacology studies where a treatment period beyond 48 hours is desirable and more accurately depicts long-term patient recovery in a clinical setting. While some neurologic deficit remained after 30 days, visible infarcts were shown to naturally resolve using TTC staining. After observing the time course of gradual improvement in both transiently and permanently occluded MCAO models, a seven-day time point was found to be most feasible for future studies using minocycline. Varying occlusion times for transiently occluded groups were also evaluated to generate the most severe reproducible infarct, and it was determined that 120 minutes was the optimal occlusion time for later transient MCAO model use.

We further used this (the 120-minute transient occlusion) and the permeant occlusion models to study the effect of minocycline treatment on infarct severity. A recent clinical study in n=152 stroke patients found that a significant neurologic improvement those receiving minocycline at 7 and 30 days, but concluded further studies were warranted.^19^ Minocycline treatment beginning post-MCAO surgery and continuing for seven days was shown to slightly improve neurologic deficit in both transiently and permanently occluded MCAO models. Direct infarct size was also linked with neurologic improvement showing a greater decrease over time in the minocycline treated groups, both permanent and transient, when compared to the control groups. Although small sample size and sample variability prevented statistical significance, a trend was evident showing a greater improvement in deficit and infarct size for the transient MCAO group as compared to the permanently occluded group, indicating the importance of revascularization for long term patient outcome.

The mechanisms of neuroprotection by minocycline are unclear. while we did not examine any mechanism for the apparent neuroprotective ability of minocycline, future studies should be performed to evaluate the most common theories: e.g., microglial activation,^9,10^ reduced matrix metalloproteinase activity,^11^ inhibition of cell death by blocking activated caspase 3 formation,^12^ or activation of the mitogen-activated protein kinase (MAPK) p38 pathway.^17^ Indeed, ED1 staining could be performed in conjunction with other proposed staining methods (GFAP-glial fibrillary acidic protein staining), to visualize activated microglia in sectioned brain slices and compare them in treatment and control groups.^17^

This study has several limitations. First, TTC staining does not allow for other histological examination of the tissue once completed. To examine the histopathology of infarcted regions (e.g., Nissl staining) additional animal groups are needed. Second, there was high variability in the resultant infarctions. This may be because the occlusions were guided only by knowledge of the rat brain vessel anatomy. Doppler flow probes could help confirm occlusion (and/or determine the percentage of occlusion that is achieved) by the suture, which then could be adjusted if needed. This could help control for inter-animal anatomical variability, which would decrease mortality and increase the consistency of the model.

## Conclusions

We implemented a permeant and transient MCAO model in female, retired breeder rats, and sacrifice them at multiple time points to determine the most appropriate time course for analysis. We then utilized the most effective models (permeant and 120-minute transient evaluated at 7 days) to evaluate neuroprotective qualities of minocycline. Here, we found that those in the treatment groups experienced a greater improvement in neurologic scores and a larger decrease in infarct size after daily minocycline treatment across all 7 days. This improvement was more prominent in the transiently occluded treatment group than in the permanently occluded group, suggesting that neuroprotectants may present a greater benefit to patients when a revascularization treatment is also provided.

