## Supplemental Figure 1 for "Evaluating Minocycline as a Neuroprotective Agent in an Aged Female Rat MCAO Stroke Model"

### Supplemental Data for: Evaluating Minocycline as a Neuroprotective Agent in an Aged Female Rat MCAO Stroke Model

University at Buffalo

Buffalo, NY 14203, USA

#### Supplemental Figures

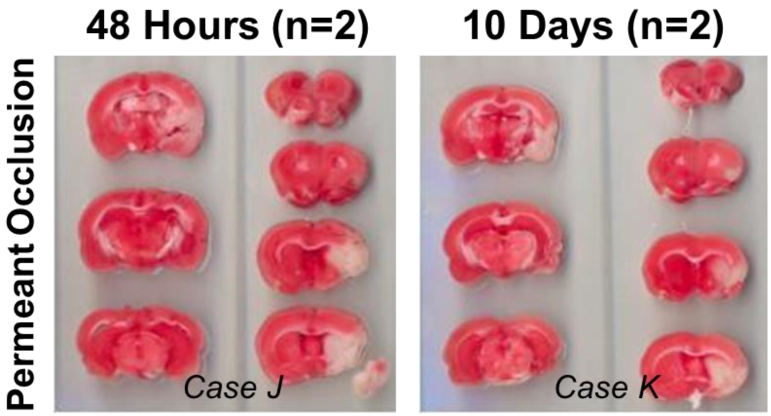


**Supplemental Figure 1: TTC staining examples of permeant occlusions that were euthanized early.** N=2 at 48 hours and n=2 at 10 days.
